# Ivermectin inhibits extracellular vesicle secretion from parasitic nematodes

**DOI:** 10.1101/2020.07.20.212290

**Authors:** Hannah J. Loghry, Wang Yuan, Mostafa Zamanian, Nicolas J. Wheeler, Timothy A. Day, Michael J. Kimber

## Abstract

Lymphatic filariasis (LF) is a disease caused by parasitic filarial nematodes that is endemic in 49 countries and affects or threatens over 890 million people. Strategies to control LF rely heavily on mass administration of anthelmintic drugs including ivermectin (IVM), a macrocyclic lactone drug considered an Essential Medicine by the WHO. However, despite its widespread use the therapeutic mode of action of IVM against filarial nematodes is not clear. We have previously reported that filarial nematodes secrete extracellular vesicles (EVs) and that their cargo has immunomodulatory properties. Here we investigate the effects of IVM and other anti-filarial drugs on parasitic nematode EV secretion, motility, and protein secretion. We show that inhibition of EV secretion was a specific property of IVM, which had consistent and significant inhibitory effects across nematode life stages and species (with the exception of male parasites). IVM inhibited EV secretion, but not parasite motility, at therapeutically relevant concentrations. Protein secretion was inhibited by IVM in the microfilariae stage, but not in any other stage tested. Our data provides evidence that inhibiting the secretion of immunomodulatory EVs by parasitic nematodes could explain, at least in part, IVM mode of action and provides a phenotype for novel drug discovery.

## 1. Introduction

Lymphatic filariasis (LF) is a Neglected Tropical Disease caused by thread-like parasitic filarial nematodes, including *Brugia malayi*, that establish in the lymphatic vasculature. LF is often asymptomatic but symptoms manifest in approximately 40% of cases with lymphedema, hydrocoele, dermatitis and long-term disability characterizing clinical disease. It is estimated that LF is endemic in 49 countries and that over 890 million people are infected or at risk of infection (World Health Organization, 2019). In 2000 the Global Programme to Eliminate Lymphatic Filariasis was created with the goal of eliminating this disease by 2020 and although there has been reduction in the prevalence of LF in some areas this disease is far from being eliminated. Strategies to control LF and other filarial parasitic nematode infections rely heavily on mass administration of the anthelmintic drugs ivermectin (IVM), albendazole (ABZ) and diethylcarbamazine (DEC) in endemic areas. IVM is classified as an essential medication by the World Health Organization (World Health Organization, 2019) and since 2000, over 7 billion treatments have been delivered to at risk individuals (World Health Organization, 2019), however, this disease still remains an issue. One challenge to eliminating LF centers on the inadequate drugs that are currently available; neither IVM, ABZ or DEC effectively kill adult parasites, thus established infections are incurable. Compounding this and despite their widespread use, the therapeutic modes of action of IVM, and to a lesser extent DEC, are not entirely clear.

A current working hypothesis for the therapeutic activity of IVM is that it inhibits the release of excretory-secretory (ES) products from parasites; this inhibition is postulated to “unmask” the parasite allowing for host recognition and parasite clearance (Moreno *et al*., 2010). In support is the acceptance that the host immune system is involved in filarial parasite elimination, especially in the clearance of microfilaria (mf) stage worms (Carithers, 2017; Wolstenholme *et al*., 2016) and data from experiments such as Berrafato *et al.* showing that IVM enhanced leukocyte binding to *Dirofilaria immitis* mf and Semnani *et al*. who showed that IVM could reverse the modified Th2 phenotypes caused in filaria infected patients (Berrafato *et al*., 2019; Semnani *et al*., 2006). There is widespread support that ES products from filarial nematodes do modulate host immune responses. Early filarial nematode infection elicits a canonical Th2 immune response characterized by increased production of the cytokines interleukin (IL)-4, IL-5, IL-9, IL-10, and IL-13 and the antibody isotypes IgG1, IgG4 (in humans), and IgE and increased production of Th2 cells, eosinophils, alternatively activated macrophages, and innate lymphoid cells 2 (ILC2) (Allen & Maizels, 2011; Geary *et al*., 2010). With development of chronic filarial infection the Th2 response becomes “modified” to a more tolerant, regulatory environment with increased IL-4, IL-10, T_reg_ and alternatively activated macrophage proliferation and reduction in IL-5, IL-13 and T cell proliferation coupled with T cell anergy and decreased antigen presenting capabilities (Babu & Nutman, 2014). There is considerable evidence that filarial nematode parasites contribute to this “modified” phenotype but the exact parasite factors driving this manipulation remain uncertain.

Extracellular vesicles (EVs) are membrane-bound vesicles secreted into the extracellular environment by eukaryotic and prokaryotic cells. Although once thought to be carriers of waste products, it has been shown that EVs function in many physiological processes and are important mediators of cell-to-cell signaling (Bobrie *et al*., 2011; Lee *et al*., 2012; Raposo *et al*., 1996; Valadi *et al*., 2007). EVs are considered a heterogenous group of sub-cellular structures that can be subdivided based on size and biogenesis. Primary focus has been on two subsets of EVs, microvesicles that range from 150-1,500 nm and a smaller grouping (30-150 nm) originally termed exosomes (Johnstone *et al*., 1987). Exosomes are products of the endosomal pathway and are derived from multivesicular bodies (MVBs) that fuse with the cell membrane to secrete the vesicles into the extracellular space (Catalano & O’Driscoll, 2020; Riaz & Cheng, 2017; Vlassov *et al*., 2012). Consistent with a role in cell-to-cell communication, EVs contain diverse functional cargo that varies depending on the cellular origin of the EVs, but in general include bioactive proteins, RNA and lipids (Thery *et al*., 2002; Valadi *et al*., 2007). EV secretion from diverse parasitic nematodes has been described (Buck *et al*., 2014; Coakley *et al*., 2017; Eichenberger, Ryan, *et al*., 2018; Eichenberger, Talukder, *et al*., 2018; Gu *et al*., 2017; Hansen *et al*., 2015, 2019; Harischandra *et al*., 2018; Shears *et al*., 2018; Tritten *et al*., 2017; Tzelos *et al*., 2016; Zamanian *et al*., 2015) and the cargo of these EVs have immunomodulatory functions (Buck *et al*., 2014; Quintana *et al*., 2017; Tritten *et al*., 2016). We have previously reported that *B. malayi* secretes EVs and that their cargo has putative immunomodulatory properties (Harischandra *et al*., 2018; Zamanian *et al*., 2015). Driven by these emerging data, EVs have been advanced as a potential mechanism by which parasites modulate host immune responses (Buck *et al*., 2014; Coakley *et al*., 2017; Eichenberger, Sotillo, *et al*., 2018; Harischandra *et al*., 2018).

We propose that nematode EVs are essential for filarial nematode parasitism and hypothesize that effective anti-filarial drugs inhibit their secretion. To investigate this hypothesis, a panel of anti-filarial drugs was screened for their ability to reduce EV secretion from parasitic nematodes. We found that IVM had the most consistent inhibitory effects on EV secretion by various species and life stages of parasite. Importantly, however, IVM had insignificant effects on motility and limited effects on protein secretion at therapeutically relevant concentrations and timepoints. These observations provide insight into the mechanism of action of IVM and may support prioritizing inhibition of EV secretion as a screenable phenotype for novel anti-filarial drug development.

## 2. Materials and Methods

### 2.1. Parasite culture and maintenance

*Brugia malayi* and *B. pahangi* parasites were obtained from the NIH/NIAID Filariasis Research Reagent Resource Center (FR3) at the University of Georgia, USA. Persistent *B*. *malayi* infections at FR3 are maintained in domestic short-haired cats. To obtain adult stage *B. malayi* jirds were infected intraperitoneally with approximately 400 L3 stage parasites. 120 days post-infection jirds were necropsied to collect adult stage parasites. L3 stage *B. malayi* were obtained from dissection of anesthetized *Aedes aegypti* 14 days post-infection. Microfilaria stage *B. malayi* were obtained from a lavage of the peritoneal cavity of a euthanized gerbil. *B. pahangi* stages were obtained in the same manner as *B. malayi* with the exception that infective L3 stage parasites were collected 11 days and 16 days post-infection, respectively. The *B. malayi* parasite supply from FR3 was supplemented with parasites from TRS Labs LLC (Athens, Georgia, USA). These supplemental parasites were tested and responded to treatments in the same manner as parasites from FR3. Upon receipt at ISU, all *B. malayi* and *B. pahangi* parasites were washed several times in warmed worm culture media (RPMI with 1% HEPES, 1% L-glutamine, 0.2% Penicillin/Streptomycin, and 1% w/v glucose [all Thermo Fisher Scientific, Waltham, MA]) and then counted and cultured at 37°C with 5% CO_2_. Adult female *Ascaris suum* were collected from an abattoir in Marshalltown, Iowa, USA. These parasites were washed multiple times in warmed *Ascaris* Ringer’s Solution (13.14 mM NaCl, 9.67 mM CaCl_2_, 7.83 mM MgCl_2_, 12.09 mM C_4_H_11_NO_3_, 99.96 mM C_2_H_3_NaO_2_, 19.64 mM KCl with Gentamycin (100 μg/ml), Ciprofloxacin Hydrochloride (20 μg/ml), penicillin (10,000 units/ml), streptomycin (10,000 μg/ml), and Amphotericin B (25 μg/ml) at pH 7.87 [all Sigma-Aldrich, St Louis, MO]) and then incubated overnight at 34°C. After 24 hrs in culture the parasites were checked for visible signs of bacterial or fungal contamination; if present the parasites were discarded.

### 2.2 Drug treatments of parasites

Parasites were cultured in the presence or absence of drug to examine effects on extracellular vesicle (EV) secretion. For *B. malayi* and *B. pahangi*, 10 adult female and 10 adult males were cultured as previously described for 24 hrs in 10 ml and 3 ml culture media, respectively, in 15 ml polypropylene centrifuge tubes (Thermo Fisher Scientific). 100 L3 or 1×10^6^ microfilariae were cultured as previously described for 24 hrs in 1 ml culture media in 1.5 ml microcentrifuge tubes (Thermo Fisher Scientific). Single adult female *A. suum* were cultured in 100 ml culture media in 250 ml sterile Erlenmeyer flask for 24 hrs as previously described. Four drugs, ivermectin, albendazole, diethylcarbamazine, and levamisole (all Sigma-Aldrich) were investigated for their effects on each life stage of the parasite species. The various drugs or DMSO (vehicle control) were added to the culture media at a final concentration of 1 μM and 0.01% respectively for screening purposes. Spent media was collected after a 24 hr incubation. Additionally, drug and control treated *A. suum* and *B. pahangi* media was collected at 2, 4, 6, and 12-hr intervals to investigate the time course of the effects of the drugs. A dose curve for the effects of ivermectin on *B. malayi* were conducted in the same manner as described above with concentrations ranging from 0.1 nM – 10μM.

### 2.3 EV Isolation and Quantification

EVs were collected as previously described using differential ultracentrifugation (Harischandra *et al*., 2018; Zamanian *et al*., 2015). Media was filtered through 0.2 μm PVDF filtered syringes (GE Healthcare, Chicago, IL) or PVDF vacuum filters (Sigma-Aldrich) and centrifuged at 120,000 x *g* for 90 minutes at 4°C. The supernatant was decanted leaving approximately 1.5 ml media to ensure that the EV pellet was not disrupted. The retained media and pellet were filtered through a PVDF 0.2 μm syringe filter and centrifuged at 186,000 x *g* for a further two hrs at 4°C. Pelleted EV samples were resuspended to 500 μl in dPBS (Thermo Fisher Scientific). EV quantification and size determination were performed using nanoparticle tracking analysis (NTA; Nano-Sight LM10, Malvern Instruments, Malvern, UK).

### 2.4 Motility Analysis

The Worminator system developed and described by Marcellino *et al.* (2012) was used to quantitatively measure motility of adult filarial nematodes in microtiter plates. Microscopic parasite life stages were quantitatively analyzed by the same software, but with methods previously described by Storey *et al*. (2014). Briefly, a single adult male or female worm was cultured in one well of a standard 24-well cell culture plate (Sigma-Aldrich). For infective L3 stage worms, 10 worms were cultured per well of a 96-well plate (Corning Inc, Corning, NY). Drug or DMSO (vehicle control) was added to each well to a final concentration of 1 μM or 0.01% respectively. Worms were incubated at 37°C and 5% CO_2_ and measurements were briefly taken, at room temperature, prior to treatment, immediately after treatment (0 hrs) and at 2, 4, 6 and 24-hrs post treatment. Measurements of the effects of doses of ivermectin ranging from 0.1 nM – 10 μM on adult female *B. malayi* were conducted at 24 hrs post treatment.

### 2.5 SF21 Cell Culture

Sf21 cells (Thermo Fisher Scientific) were maintained in Ex-Cell 420 serum free media (Sigma Aldrich) with 1% Penicillin/Streptomycin and 0.25 ug/ml Amphotericin B. Cells were seeded at a density of 3×105 cells/well in a 6-well plate. After an overnight incubation at 28°C, cells were either treated with a final concentration of 1.0 μM ivermectin, 0.1 μM ivermectin or 0.01% DMSO (vehicle control). Spent media was collected after 24 hrs and processed for EV isolation as described above with the addition of an initial centrifugation step of 12,000 x *g* for 30 minutes at 4°C to eliminate cellular debris.

### 2.6 Protein Quantification Assay

A single *B. malayi* adult or 100,000 microfilariae were cultured per well of a 24-well plate with either drug or DMSO at a final concentration of 1.0 μM or 0.01%, respectively, for 24 hrs. A concentration-response curve for ivermectin was conducted on adult female worms with concentrations ranging from 0.1 nM – 10 μM. Spent media was collected and filtered through a 0.2 μM PVDF membrane filter (GE Healthcare) 500 μl of media were concentrated using a 0.5 mL, 3,000 Da Amicon Ultra centrifugal filter unit (Sigma-Aldrich) according to manufacturer’s instructions. Media samples were concentrated by centrifuging at 14,000 x *g* for 30 minutes. Samples were then washed with 500 μl dPBS for 30 minutes at 14,000 x *g*. The washing step was repeated four times. The volume of each sample was determined, and all samples were normalized to 150 μl with dPBS. Adult female samples were then further diluted 4-fold with dPBS while adult male, L3 stage and microfilariae were diluted 2-fold with dPBS to ensure that readings would fall within the standard curve. Total protein was quantified with Pierce micro BCA kit (Thermo Fisher Scientific) according to manufacturers’ instructions. Protein assay plates were quantified using a SpectraMax M2e plate reader (Molecular Devices, San Jose, CA).

### 2.7 Statistical Analysis

Due to some variation among individual parasites and between batches of parasites, experiments were conducted across multiple batches of parasite shipments, with each N representing parasites from independent shipments. Individual control and treated worms within each batch were paired together to help account for batch variation. EV NTA data was analyzed via a ratio-paired T-test with p-values less than 0.05 being considered significant. Non-linear regressions with least squares fit were used to analyze the dose response curves for ivermectin on *B. malayi* adult female EV secretion, motility, and protein secretion. Motility data was analyzed via a RM 2-way ANOVA with Geisser-Greenhouse correction followed by a Dunnet’s multiple comparison test. Paired T-test between each treatment for each life stage were used to analyze data from the protein assay as the data contained values unsuited for a ratio-paired T-test. Due to the variability among batches and individual parasites the ROUT outlier identification method was used to identify outliers in the data (Q = 0.5%). All statistical analyses were performed using Prism 8.4.1. (GraphPad Software, San Diego, CA).

## 3. Results

### 3.1 Ivermectin inhibits EV secretion from Brugia malayi in a sex- and stage-specific manner

In this study we investigated the effects of ivermectin (IVM) on *B. malayi* EV secretion *in vitro*. Parasites were cultured at 37°C in the presence or absence of IVM and the number of EVs secreted by the worms was quantified by nanoparticle tracking analysis (NTA). An initial screening concentration of 1.0 μM IVM significantly reduced EV secretion after 24 hrs incubation from *B. malayi* adult females by 59% (p = 0.0204, N = 13) and from L3 stage parasites by 31% (p = 0.0067, N = 19). There was no significant effect on EV secretion from adult male (p = 0.4028, N = 10) or microfilariae (mf) (p = 0.2081, N = 13) life stages (Fig 1A-D). The IVM concentration response in adult female *B. malayi* was further profiled and the IC_50_ determined to be 7.7 nM (Fig 1E). Studies conducted on the pharmacokinetics of a single dose of IVM in human subjects have determined serum levels to be between 20-70 nM (González Canga *et al*., 2008). IVM therefore inhibits EV secretion in adult female parasites at therapeutically relevant concentrations suggesting that this phenomenon may contribute to IVM therapeutic mode of action.

**Figure 1.**
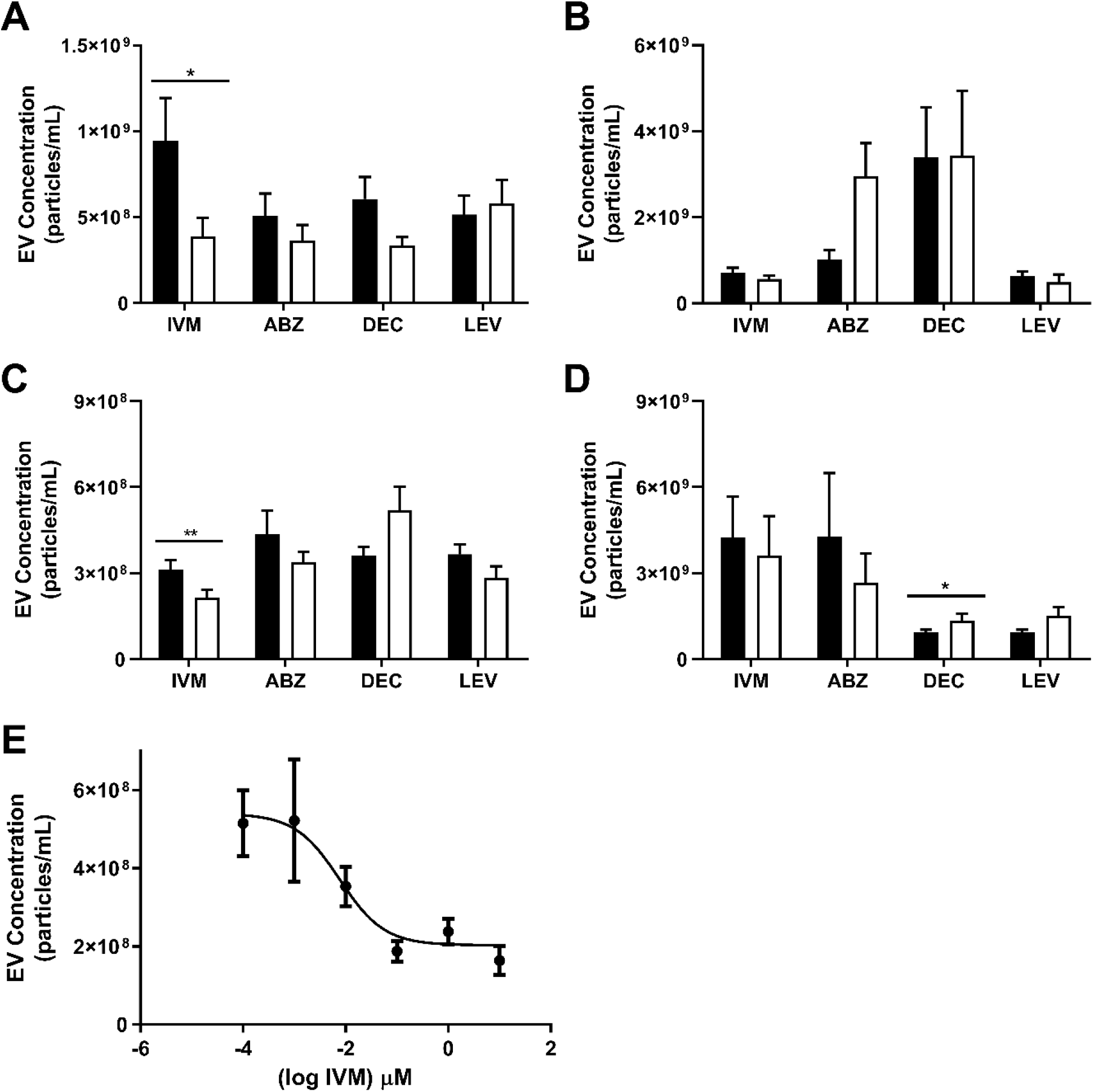
Ivermectin inhibits Brugia malayi EV secretion in a stage- and sex-specific manner. *B. malayi* life stages were cultured at 37°C in RPMI with either drug (1.0 μM) or DMSO (vehicle control). Media was collected after 24 hrs and EVs were isolated and quantified. 1.0 μM IVM significantly reduced EV secretion from adult female worms (A) and L3 stage parasites (C) but not from adult males (B) or microfilaria (D). Albendazole (ABZ), diethylcarbamazine (DEC) or levamisole (LEV) had no effect on EV secretion from any life stage except in microfilaria, where DEC increased EV secretion. (E) The IC_50_ for IVM on adult female worms was determined to be 7.7 nM. N = 7 (minimum), Mean ± SEM, *P<0.05, **P<0.01. ■ = DMSO control, □ = Treatment

These observations on the inhibitory effect of IVM on EV secretion from adult female and L3 stage *B. malayi* generally align with preliminary data previously reported, (Harischandra *et al*., 2018) with the exception of the lack of inhibition in adult males and a reduced inhibition in mf stages. The previously reported inhibitory effect on EV secretion from adult males was marginal, but the lack of effect on mf is more surprising considering its prior robustness. Previously, mf were incubated with IVM at ambient temperature whereas here they were incubated at 37°C. To test the impact of temperature on the IVM phenotype in mf, we repeated the mf IVM incubation at ambient temperature. Unlike at 37°C, 1.0 μM IVM significantly inhibited EV secretion by 46% (p = 0.0177, N = 8) at ambient temperature (22°C) (Fig 2A). Control and treated parasites were still viable at the end of the experiment indicating that it was not loss of viability or death of the parasites that had caused inhibition of EV secretion. There are clear temperature-dependent effects on EV secretion from mf stage worms, not only did incubation of mf at 37°C abrogate the inhibitory effect of IVM on EV secretion, but it also increased EV secretion in untreated worms by approximately a factor of 7. Whilst logical to assume temperature changes impact worm physiology, there is a lack of data on the effects of temperature on specific processes and functions in mf stage nematodes. We do know that host temperature has no effect on the nocturnal periodicity of mf (Hawking, 1967) or on the ability of mf to bind to vascular endothelial cells (Schroeder *et al*., 2017). Environmental temperature may affect other physiological processes in mf leading to this increased production of EVs, but further investigation into this phenomenon is necessary.

**Figure 2.**
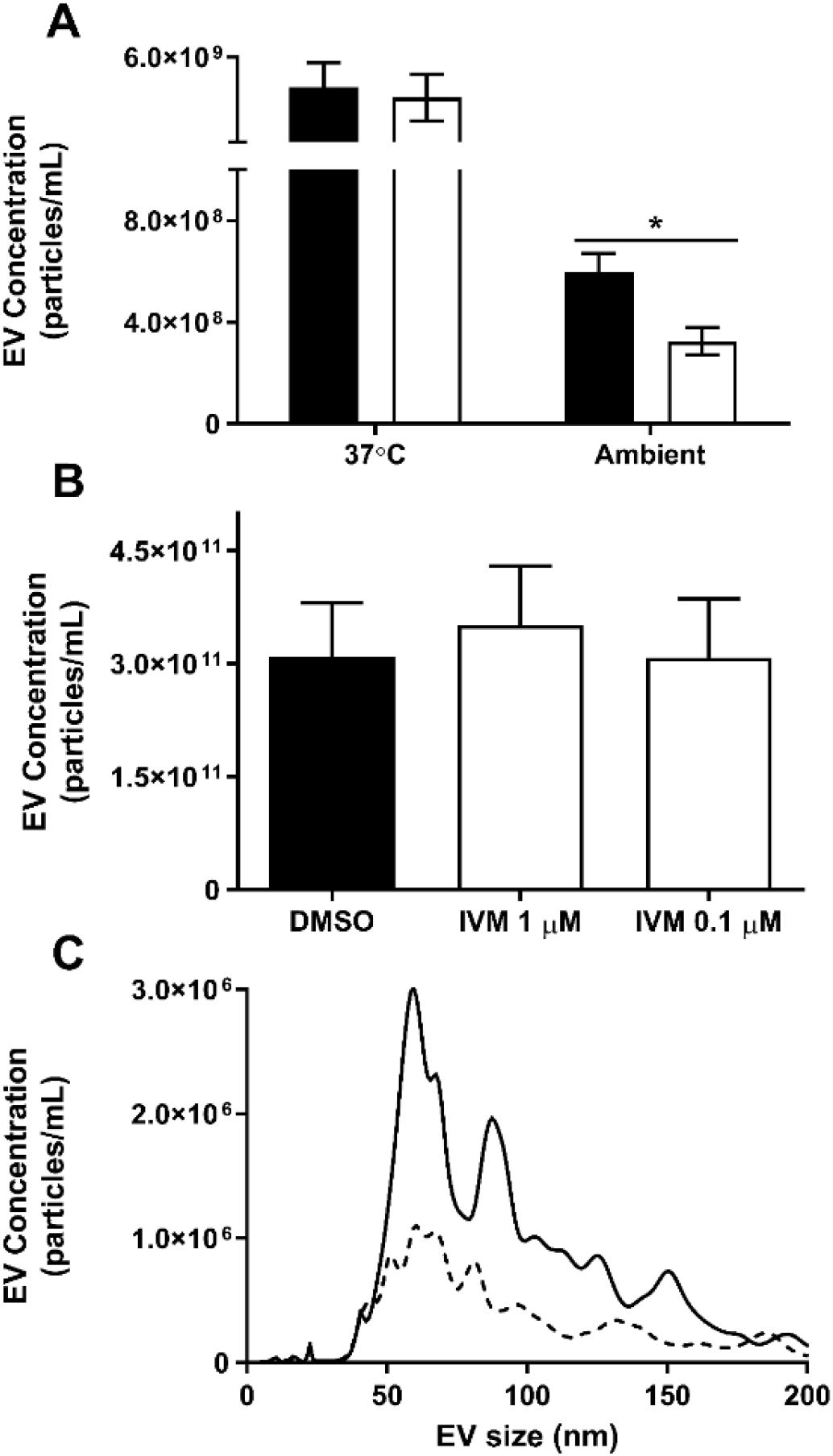
Ivermectin activity against mf stage worms is temperature dependent and does not alter EV biogenesis. *B. malayi* life stages were cultured at 37°C in RPMI with either drug (1.0 μM) or DMSO (vehicle control). Media was collected after 24 hrs and EVs were isolated and quantified. The activity of IVM on *B. malayi* microfilariae was temperature dependent (A). Inhibition of EV secretion occurred at ambient temperature, but not at 37°C. To investigate IVM mechanism of inhibition Sf21 cells were treated with either DMSO, high (1.0 μM) or low (0.1μM) concentrations of IVM. Spent media was collected after 24 hrs and EVs were collected and quantified. (B) there were no differences in the ability of DMSO control, high, or low concentrations of IVM treated cells to secrete EVs. To further investigate the potential effect on biogenesis the size of EVs secreted from adult female parasites were evaluated (C). The mean size of EVs secreted from adult females was highly similar between DMSO and IVM treated parasites. N = 3 (minimum). *P<0.05. ■ = DMSO, □ = Treatment. Solid line = DMSO, Dashed line = IVM.

This temperature-dependent effect in mf prompted further efforts to determine the mechanism by which IVM is inhibiting EV secretion. One hypothesis is IVM inhibits EV biogenesis in these parasites. Nematode cell lines that might provide an opportunity to interrogate EV biogenesis are lacking so Sf21cells from the army fallworm, *Spodoptera frugiperda*, were used as an ecdysozoan surrogate to examine the effect of IVM on EV biogenesis *in vitro*. Sf21 cells were treated with high (1.0 μM) and low (0.1 μM) concentrations of IVM and EV secretion was analyzed by NTA after 24 hrs. Neither concentration of IVM inhibited EV secretion from Sf21 cells (Fig 2B), perhaps suggesting the drug does not directly impact EV biogenesis. Another hypothesis is that IVM disrupts the location from where EVs are secreted. The physical characteristics of EVs produced from IVM-treated or DMSO-treated *B. malayi* adult females were analyzed by NTA. Although EVs are, as a whole, a heterogenous population in regards to size, our data suggested that there was no obvious shift in EV size profile following IVM treatment (Fig 2C). It might be expected that disrupting the location from where EVs are secreted, for example the parasite GI tract or reproductive structures, might alter their physical characteristics, but this was not observed. The physical characteristics of EVs secreted by other *B. malayi* life stages were also analyzed (Supplemental Figure 2A-C). Consistent with the adult female data, there were no differences in EV size following IVM treatment of any life stage. Collectively, our data suggests that IVM works not by altering EV biogenesis or changing the location from where EVs are secreted, but rather restricts the numbers of EVs being secreted by pre-existing pathways.

### 3.2 Other drugs with anti-filarial activity do not inhibit EV secretion from *B. malayi*

To examine if inhibition of EV secretion is a general feature of drugs with anti-filarial activity, we tested a panel of drugs with known anti-filarial activity including albendazole (ABZ), diethylcarbamazine (DEC) and levamisole (LEV). LEV is a nicotinic agonist and although more typically used to treat gastrointestinal nematode infection, was included in the panel because it is an anthelmintic drug with known neuromuscular effects on filarial nematodes (Martin, 1997; Robertson *et al*., 2013) and also because of reported microfilaricidal effects on canine heartworm, *D. immitis* (Carlisle *et al*., 1984; Mills & Amis, 1975). Parasites were cultured with or without an initial screening concentration of 1.0 μM drug and EVs were quantified using NTA. 1.0 μM DEC significantly increased EV secretion from *B. malayi* mf by 43% (p = 0.0177, N = 7) after 24 hrs (Fig 1D). DEC also seemed to increase EV secretion in L3 (Fig 1C) though not significantly (p = 0.2236, N = 10). Additionally, DEC had a moderate, but not significant, inhibition on EV secretion in adult females (p = 0.1704, N = 10) and had no effect on adult males (p = 0.2323, N = 9) (Fig 1A-B). There was a minor, but not significant, inhibition of EV secretion due to ABZ on *B. malayi* adult females (p = 0.4042, N = 10), L3 (p = 0.1564, N = 11), and mf (p = 0.7815, N = 7) (Fig 1A,C,D). In contrast, ABZ treatment seemed to increase EV secretion from adult males (Fig 1B) though not significantly (p = 0.0821, N = 9). LEV did not have an effect on EV secretion from either male (p = 0.1091, N = 11) or female adults (p = 0.8659, N = 9) (Fig 1A-B). L3 stage parasites showed a minor although not significant inhibition in EV secretion due to LEV treatment (p = 0.0719, N = 10) while mf had a moderate, but not significant increase in EV secretion due to LEV treatment (p = 0.0770, N= 7) (Fig 1C-D). In summary, none of the drugs in the panel significantly inhibited EV secretion from any of the *B. malayi* life stages when tested at 1.0 μM, except for DEC which significantly increased EV secretion from *B. malayi* microfilariae. This is significant because it suggests that inhibition of EV secretion by filarial nematodes may be a phenotype specific to IVM treatment and is not observed upon treatment with other anthelmintic drugs that are known to have anti-filarial activity, including a drug (LEV) that has clear neuromuscular effects on filarial worms.

This helps support the hypothesis that the mechanism of action of IVM, and perhaps other macrocyclic lactones, includes inhibition of EV secretion.

### 3.3 Ivermectin has broad inhibitory effects on EV secretion across other filarial and gastrointestinal nematode parasites

To test whether the inhibitory IVM phenotype in *B. malayi* was broadly consistent in nematodes, we repeated the same screening experiment with our same drug panel using first a related species of filarial nematode, *B. pahangi*. Our analysis was limited to *B. pahangi* adult females, adult males and mf based on worm availability. All drugs in the panel significantly inhibited EV secretion from *B. pahangi* adult female parasites. IVM treatment had the greatest reduction in EV secretion with an inhibition of 63% (p = 0.0083, N = 10), while ABZ had an inhibition of 61% (p = 0.0066, N = 12), DEC by 59% (p = 0.0322, N = 12) and LEV by 44% (p = 0.0416, N = 11) (Fig 3A). The IVM, ABZ and DEC results generally paralleled the trends seen in *B. malayi* adult females, but with LEV now also active. Neither IVM (p = 0.9897, N = 11), ABZ (p = 0.9369, N = 13), DEC (p = 0.0925, N = 12), nor LEV (p = 0.7558, N = 11) had any effect on adult male *B. pahangi* (Fig 3B). Again, this is consistent with the results seen in *B. malayi* adult male parasites. In mf stage *B. pahangi*, IVM significantly reduced EV secretion by 40% at 37°C (p = 0.0358, N = 4) (Fig 3C), which contrasts sharply with what was seen in *B. malayi* mf and perhaps better aligns with the expected bioactivity of IVM at this life stage (Moreno *et al*., 2010). While DEC significantly increased EV secretion in *B. malayi* mf it did not have any effect on *B. pahangi* mf (p = 0.6428, N = 3) (Fig 3C). Lastly, ABZ (p = 0.6605, N= 3) and LEV (p = 0.4125, N = 3) had no effect on EV secretion from *B. pahangi* mf (Fig 3C). To investigate whether the inhibitory effects of IVM on EV secretion were seen in more divergent nematode species we again repeated our screen on single adult female *Ascaris suum*, a soil-transmitted gastrointestinal nematode. IVM (p = 0.0013, N = 14) and LEV (p = 0.0021, N = 16) both significantly inhibited EV secretion from individual adult female *A. suum* after 24 hrs by 99.4% and 99.1% respectively (Fig 3E). However, neither ABZ (p = 0.3769, N = 12) nor DEC (p = 0.9680, N = 16) had any effect on EV secretion (Fig 3E).

**Figure 3.**
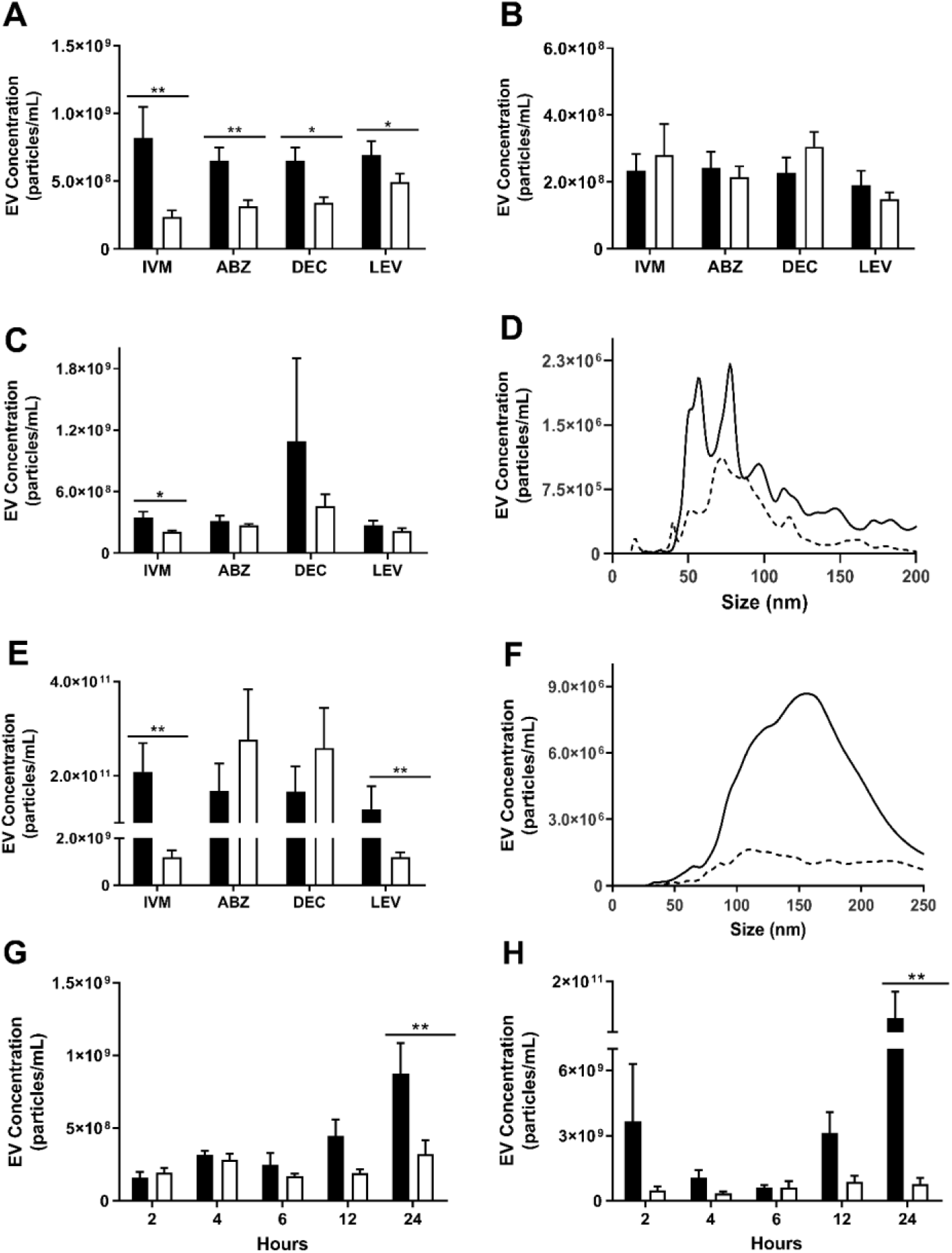
Ivermectin has broad inhibitory effects across filarial and gastrointestinal parasites. *B. pahangi* life stages in RPMI and adult female *A. suum* in *Ascaris* Ringers solution were cultured at 37°C with either drug (1.0 uM) or DMSO (vehicle control). Media was collected after 24 hrs and EVs were isolated and quantified. 1.0 μM IVM, ABZ, DEC, and LEV all significantly inhibited EV secretion from *B. pahangi* adult female parasites (A) while only IVM significantly reduced EV secretion in the microfilariae life stage (C). No treatment had any effect on EV secretion from *B. pahangi* adult male parasites (B). For *A. suum* adult females, both IVM and LEV significantly inhibited EV secretion (E). The size distribution of EVs secreted from DMSO and IVM treated *B. pahangi* adult females (D) and *A. suum* adult females (F) were highly similar indicating that IVM does not affect the physical characteristics of EVs produced (F). IVM rapidly inhibits EV secretion from adult female *B. pahangi* and *A. suum* 24 hrs post-treatment (G-H). N = 3 (minimum). Mean ± SEM, *P<0.05, **P<0.01. ■ = DMSO, □ = Treatment. Solid line = DMSO, Dashed line = IVM.

The size characteristics of those EVs that continued to be secreted by both adult female *B. pahangi* (Fig 3D) and *A. suum* (Fig 3F) following IVM treatment was examined using NTA. As was seen in *B. malayi*, IVM did not affect the physical characteristics of EVs produced from these other species of parasites, providing additional evidence that the inhibitory bioactivity of IVM may not be involved in changing from where EVs are secreted. The effects of the other drugs in our screening panel on the size of EVs secreted by *B. malayi* and *B. pahangi* life stages were also analyzed (Supplemental Fig 2A-E). None of the drugs tested had any effect on the physical characteristics of EVs secreted.

*B. pahangi* and *A. suum* adult female worms secrete EVs more robustly than *B. malayi.* In the case of *A.suum*, they secrete approximately 250 times more EVs than *B. malayi* in 24 hrs. This positioned us to use these two species to better understand how rapidly IVM inhibits EV secretion from susceptible parasitic nematodes. Adult female *B. pahangi* and *A.suum* parasites were treated with 1.0 μM IVM as before and spent media collected at 2, 4, 6, 12 and 24 hrs post IVM treatment. IVM significantly inhibited both *B. pahangi* and *A. suum* EV secretion by 63% (p = 0.0076, N = 3) and 99.4% (p = 0.0054, N = 3), respectively, at the 24 hrs post-treatment timepoint (Fig 3G-H). In addition, EV secretion was inhibited as early as at 12 hours post-treatment by 72% (p = 0.1929, N = 3) for *A. suum* and by 57% (p = 0.0762, N = 3) for *B. pahangi*, though not statistically significant (Fig 3G-H). *In vivo* studies have shown IVM reduces microfilaremia in mice experimentally infected with *B. malayi* 24 hrs post-treatment (Halliday *et al*., 2014). Thus, the rapidity of onset for this EV secretion phenotype is consistent with the therapeutic action of IVM. For context, other IVM phenotypes have been identified at 24 hrs post-treatment timepoint, *in vitro* IVM reduces the release of *B. malayi* mf from adult female worms (Tompkins *et al*., 2010) and increases the binding of polymorphonuclear leukocytes to *B. malayi* mf (Berrafato *et al*., 2019) after 24 hrs. In canine heartworms, IVM inhibits the motility of Missouri strain *D. immitis* 24 hrs post-treatment *in vitro* (Maclean *et al*., 2017).

### 3.4 Ivermectin inhibition of EV secretion is not driven by loss of gross motor function

Glutamate-gated chloride channels (GluCl) and nicotinic acetylcholine receptors (nAChRs), are known targets for IVM and LEV, respectively (Arena *et al*., 1991, 1992; Harrow & Gration, 1985). GluCl have been identified in motor neurons and interneurons in various parasitic nematode species (Adrian J. Wolstenholme & Rogers, 2005) and nAChRs have been identified at the neuromuscular junction in filarial nematodes (Martin, 1997). Their locations lead to the discovery that IVM can induce paralysis of pharyngeal pumping in *Haemonchus contortus* (Geary et al., 1993) and both IVM and LEV can cause paralysis of *B. malayi* parasites (Mostafa *et al*., 2015; Tompkins *et al*., 2010). Due to these documented effects it is plausible that the EV phenotype is driven by gross motor function defects. To test this we examined the effect of our screening panel on gross motor function by analyzing motility quantified using the Worminator software system (Marcellino *et al*., 2012). Our analysis was limited to *B. malayi* adults and L3 stage parasites due to parasite availability and difficulties in consistently recording the smaller mf life stage. A single *B. malayi* adult or 10 L3 stage parasites were cultured in a 24-well or 96-well plate respectively with or without 1.0 μM drug. Video recordings were taken prior to treatment, immediately after treatment (0 hrs), and at 2, 4, 6 and 24 hrs post-treatment. IVM significantly reduced adult female motility by 57% beginning at 4 hrs post-treatment (p < 0.0001, N=5) (Fig 4A). However, when the kinetics of IVM treatment on adult female parasites was investigated it was observed that more therapeutically relevant concentrations did not affect motility (Fig 4D). This is corroborated by other data that shows that motility of *B. malayi* parasites was not inhibited by concentrations of IVM less than 2 μM (Storey *et al*., 2014; Tompkins *et al*., 2010). This provides additional evidence that therapeutically relevant concentrations of IVM do not affect filarial nematode motility. The IC_50_ for IVM was determined to be 0.203 μM. The IC_50_ for EV secretion (7.7 nM) was below that of motility indicating that IVM is not reducing EV secretion by paralyzing the parasites. DEC (p < 0.001, N=5) significantly inhibited adult female motility immediately upon treatment, but parasites began to recover at one hr (p < 0.01, N=5) and completely recovered by 4 hrs post-treatment. LEV (p < 0.001, N=5) significantly inhibited adult female motility by 88% immediately upon treatment, but parasites began to recover during the remaining 24 hrs. At one hr post treatment LEV significantly inhibited adult female motility by 71%, at four hrs by 40%, at six hrs by 35% and at 24 hrs by 29% (1-6 hrs post-treatment: p < 0.0001, N=5; 24 hrs: P < 0.01, N = 5). As was discussed earlier, no drug in our panel had any effect on EV secretion in adult male *B. malayi,* but it was discovered that LEV was a potent inhibitor of motility in adult male *B. malayi* (Fig 4B). Motility in adult males was significantly inhibited by 70% upon treatment with LEV. Adult male parasites treated with LEV did not recover with significant inhibition ranging from 70-76% over the 24 hrs tested (1-24 hrs post treatment: p < 0.0001, N=5). The only drug that had any effect on *B. malayi* L3 parasites was also LEV with inhibition of motility by 90% immediately upon treatment (p < 0.001, N=5) (Fig 4C). However, a very quick recovery of motility was seen in just one hour. In summary, 1.0 μM IVM had inhibitory effects on *B. malayi* adult female motility, but this concentration does not compare to therapeutically relevant concentrations or to the concentrations that inhibited EV secretion. LEV also had inhibitory effects on motility in all life stages tested, but adult female and L3 life stages recovered over 24 hrs while adult males did not recover. Due to the differences in IVM effects on motility compared to EV secretion we can conclude that inhibition of EV secretion is not a factor of parasite gross motor function being compromised.

**Figure 4.**
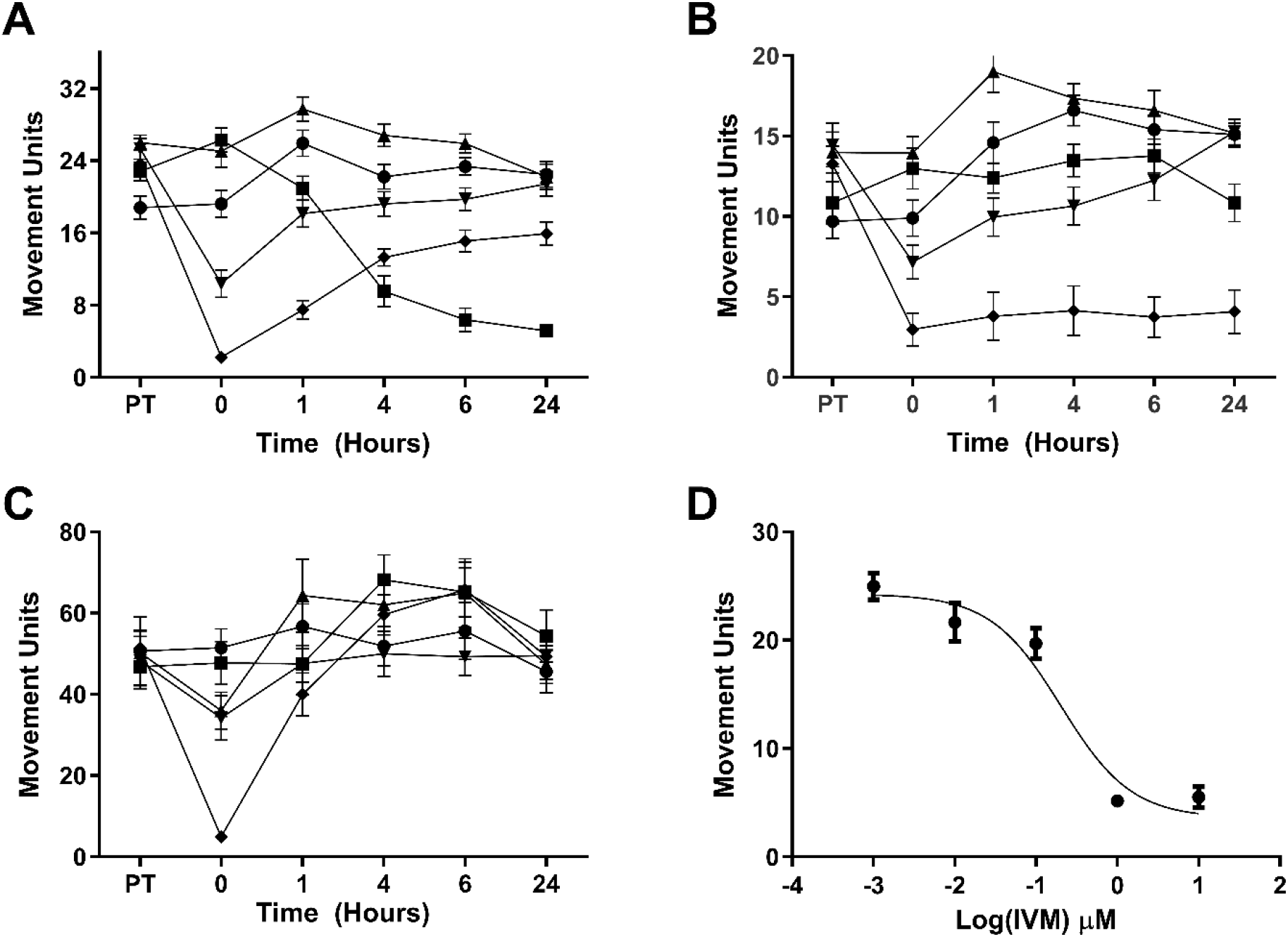
Ivermectin inhibition of EV secretion is not driven by loss of gross motor function. Single adult or 10 L3 stage *Brugia malayi* parasites were cultured in a 24-well plate with either 1.0 μM drug or DMSO (vehicle control). Video recordings of worms were taken at timepoints ranging from pre-treatment to 24 hrs using the Worminator system. (A) IVM significantly reduced adult female *B. malayi* motility as compared to control from 4-24 hrs post treatment (p<0.0001). However, further investigation revealed that more therapeutically relevant concentrations of IVM did not affect adult female motility. The IC_50_ for IVM on adult female parasites was determined to be 203 nM. (D). DEC significantly reduced adult female motility immediately upon treatment and one hour post treatment (p<0.001). LEV significantly reduced adult female motility from 0-24 hrs (p<0.0001-p<0.01) though the parasites began to recover after initial treatment. (B) DEC significantly reduced adult male motility from one to four hours post treatment (p<0.05, p<0.01) and IVM began to reduce motility at 24 hrs post treatment (p<0.05). LEV significantly reduced motility with no recovery from 0-24 hrs post treatment (p<0.0001). (C) LEV significantly reduced L3 stage motility immediately upon treatment (p<0.0001), but the parasites quickly recovered within one hr. N = 11 (minimum), Mean ± SEM. ● = DMSO, ■ = IVM, ▲ = ABZ, ▼ = DEC, ♦ = LEV

### 3.5 Ivermectin does not have parallel effects on EV and protein secretion

Data indicate that IVM and ABZ inhibit protein secretion from *B. malayi* mf (Moreno *et al*., 2010). We have already shown that IVM can inhibit EV secretion from *B. malayi* so we hypothesized that excretory-secretory (ES) proteins and EVs would similarly be affected by the screening drug panel. Parasites were cultured with or without drug and ES protein secreted into the culture media was quantified 24 hrs after treatment using BCA. We chose to examine the 24 hr time point as it is the most consistent with IVM therapeutic mechanism of action. Unlike EV secretion, neither 1.0 μM IVM (p = 0.8152, N = 5), ABZ (p = 0.9962, N = 5), DEC (p = 0.8863, N=5) or LEV (p = 0.1571, N = 5) had any effect on protein secretion from *B. malayi* adult females (Fig 5A,D). Similarly, no effect of any drug on protein secretion was observed in adult males (IVM p = 0.9400, N = 5, ABZ p = 0.4906, N=5, DEC p = 0.9902, N = 5, LEV p = 0.8043, N = 5) (Fig 5B). Inhibition of ES protein secretion from *B. malayi* mf was noted after treatment with 1.0 μM IVM (23%, p = 0.0535, N = 5) (Fig 5C), however, ABZ had no effect on protein secretion (p = 0.5827, N = 5). DEC (0.5789, N=5) and LEV (p = 0.1648, N = 5) also had no inhibitory effect on ES protein secretion from *B. malayi* mf. The data do not exactly correlate to previously published work describing clear inhibitory effects of IVM and ABZ protein release from *B. malayi* mf (Moreno *et al*., 2010). The assay we used to quantify ES protein secretion from *B. malayi* was slightly modified from that study but was fundamentally the same. Despite tight technical replication, there was challenging biological variability between worm batches and the low quantities of ES protein secreted necessitated a concentration step, potentially exacerbating variability. Despite this, we have high confidence in comparing this data with those of Moreno *et al* (2010). Both studies observed a rapid inhibition of ES protein secretion from *B. malayi* mf following 24 hr treatment with 1.0 μM IVM, although our observed inhibition was moderately higher at 23% inhibition than Moreno *et al* with 14%. Further, Moreno *et al* described an inhibitory effect of ABZ (more potent than that of IVM) that was not observed here.

**Figure 5.**
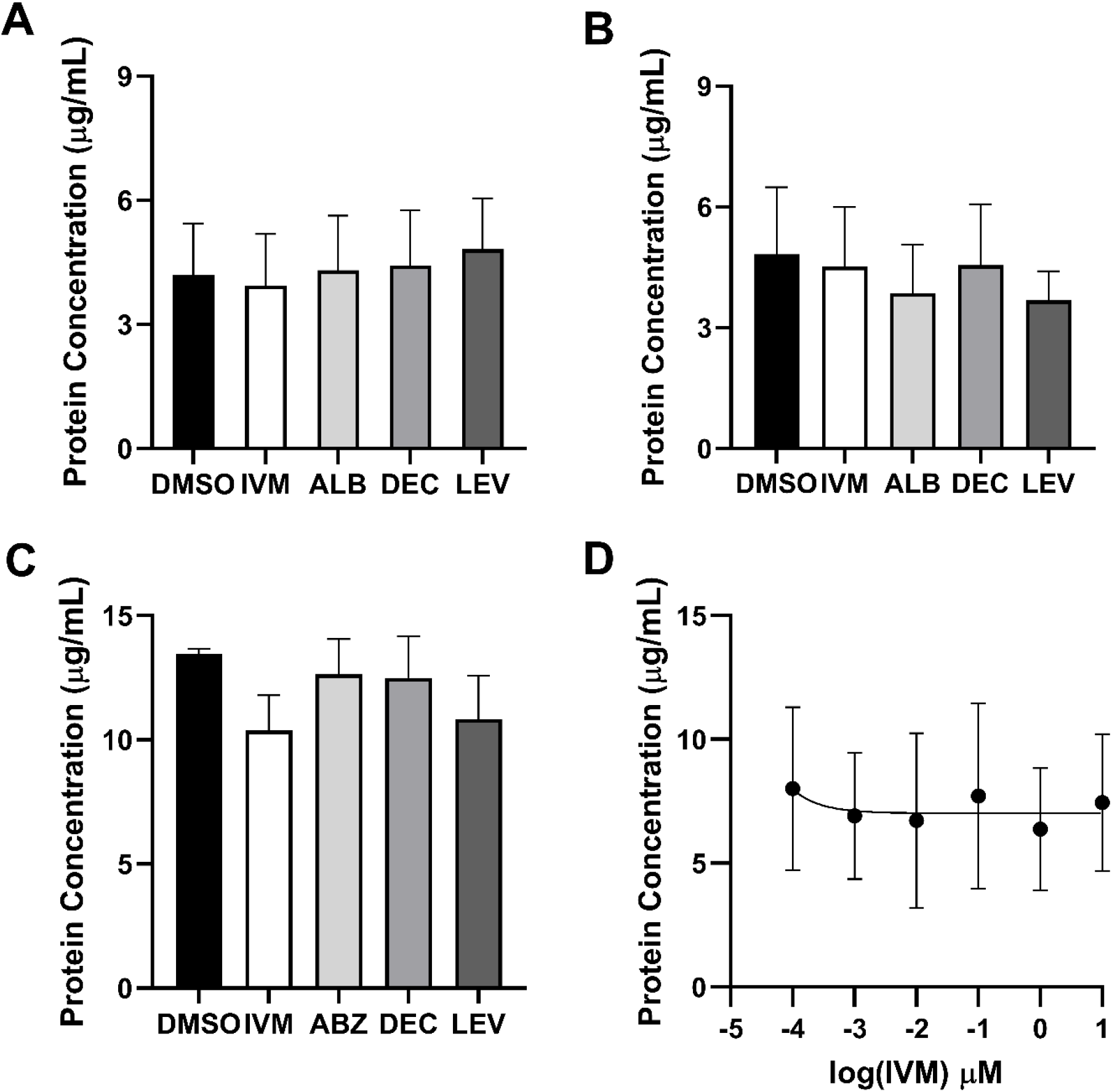
EV and protein secretion are differentially affected by ivermectin. Single adult or 100,000 microfilariae *Brugia malayi* parasites were cultured per well of a 24-well plate with either drug (1.0 μM) or DMSO (0.01%) for 24 hrs. Spent media was collected and proteins were concentrated and washed. Protein concentration was determined by absorbance at 562 nm. No drug had any effect on protein secretion from *B. malayi* adult females (A). *B. malayi* adult males (B) had a decrease in protein secretion due to LEV though not statistically significant. Microfilariae protein secretion (C) was inhibited by ivermectin (p = 0.0535). (D) A dose response curve for IVM on adult female parasites showed no effect from any concentration of IVM (10 μM – 0.1 nM). N = 5 (minimum), Mean ± SEM.

## 4. Discussion

IVM is a broad spectrum, anti-parasitic drug that is commonly used to treat and prevent multiple diseases caused by parasitic nematodes. Even with its extensive usage, the therapeutic mechanism of action of this drug is not completely understood (Wolstenholme *et al*., 2016). The current accepted hypothesis is that it functions, at least in part, by inhibiting secretion of ES proteins from parasites thereby “unmasking” the parasites and facilitating host clearance (Moreno *et al*., 2010). Recent work has led to the characterization of the *B. malayi* secretome which, combined with RNA sequencing approaches, has defined a complex milieu of proteins and miRNAs secreted from these and other filarial parasites (Bennuru *et al*., 2009; Hewitson *et al*., 2008; Hoy *et al*., 2014; Kaushal *et al*., 1982; Tritten *et al*., 2016). Within this heterogenous mix of ES products are documented host immunomodulatory effectors, including leucyl aminopeptidase (ES-62) and macrophage inhibiting factor 1 (MIF-1), among others (Harnett *et al*., 1998; Lal *et al*., 1990; Pastrana *et al*., 1998). Therefore, the observation that IVM can inhibit the secretion of immunomodulatory ES products is consistent with the rapid mf clearance observed after treatment in infected individuals. Although there is evidence tying the inhibition of ES product secretion to the mode of action of IVM, the critical ES products being inhibited are not immediately clear. In addition to freely secreted proteins, we have identified that prodigious numbers of EVs are also found in the ES products of filarial nematodes (Zamanian *et al*., 2015). In this study we show that IVM significantly and consistently inhibits EV secretion from *B. malayi* adult female, L3 and mf life stages, from *B. pahangi adult* females and mfs, and from female gastrointestinal *A. suum* nematodes. This inhibition occurs at therapeutically relevant concentrations (IC_50_ = 7.7 nM in adult female *B. malayi*) and time frame (within 24 hrs and perhaps even by 12 hrs). Given these properties, it is reasonable to hypothesize that the therapeutic mechanism of action of IVM against filarial nematodes may, in part, involve inhibition of EV secretion. Although premature, there is growing evidence to support this hypothesis. First, filarial nematode EVs are discrete structures that are enriched in immunomodulatory molecules. The cargo of *B. malayi* EVs includes proteins and miRNAs that have immunomodulatory functions and include modulatory proteins such as galectins and MIF-1 as well as miRNAs with identity to immunomodulatory host miRNAs (Harischandra *et al*., 2018; Zamanian *et al*., 2015). EVs from other nematode species are similarly imbued; protein and small RNA profiling of EV cargo from a range of gastrointestinal and filarial nematodes reveals a multitude of putative effector molecules with emerging functionality at the host-parasite interface (Buck *et al*., 2014; Eichenberger, Ryan, *et al*., 2018; Eichenberger, Talukder, *et al*., 2018; Gu *et al*., 2017; Hansen *et al*., 2015, 2019; Shears *et al*., 2018; Tritten *et al*., 2017). It is reasonable to posit that specifically inhibiting EV secretion would obstruct the immunomodulatory capabilities of these parasites. Second, the pharmacological distruption of EV secretion does not perfectly correlate with the secretion of other ES products, hinting that the regulation of EV secretion may be distinct to that of other ES products and therefore differentially “druggable”. IVM (1.0 μM) inhibited EV secretion from mf stage *B. malayi* and *B. pahangi* after 24 hrs by 46% and 40%, respectively. In comparison, we found the same treatment inhibited protein secretion by *B. malayi* mf more modestly at 23%. Further, whilst IVM (1.0 μM) inhibited EV secretion in adult female *B. malayi* by 59% after 24 hrs, the same treatment had no significant effect on protein secretion from those worms. Clearly, more work is needed to understand how parasite secretions are regulated but moving forward it may be advisable to disentangle the broad panoply of ES products and investigate them individually to help better understand host-parasite interactions and particularly how drugs affect the secretion of parasite effector molecules.

Parasite motility has long been used as an assay to identify and measure the anthelmintic activity of compounds and as marker of parasite health and viability. Impaired motility alone, however, does not adequately account for the therapeutic effects of IVM in filarial disease. The *in vitro* IVM concentrations that are required to produce detrimental effects on gross filarial nematode motility are significantly higher than the bioavailable concentrations found *in vivo* after therapeutic administration (González Canga *et al*., 2008; Marcellino *et al*., 2012; Storey *et al*., 2014; Tompkins *et al*., 2010). Our data support this – the IC_50_ of IVM in the *B. malayi* adult female EV secretion assay was 7.7 nM but was over 200 nM for the motility assay. IVM inhibits EV secretion but not motility in key stages at therapeutically relevant concentrations, supporting inhibition of EV secretion as a component of IVM mode of action. IVM also exerted inhibitory effects on *B. malayi* adult females but not adult males, and larval stages. This stage- and sex-specific activity does correlate to the expression patterns of genes encoding subunits of glutamate-gated chloride channels (GluCls), a proposed target for IVM (Arena *et al*., 1991, 1992). Li *et al* (2014) found that *avr-14*, a gene encoding a putative GluCl subunit in *B. malayi*, was expressed in both female and male reproductive tissues but consistently more strongly in female tissues (ovaries and surrounding body wall muscle) than male. This differential expression profile may help explain why EV secretion was inhibited in female worms but not male worms and may also point to reproductive structures as a source of these EVs in adult female worms. Proteomic analyses of EV cargo has proved valuable in identifying markers of tissue origin in other nematodes (Buck *et al*., 2014) but our previous nano-scale proteome profiling of *B. malayi* female and male EVs did not identify any clear markers supporting a reproductive tissue origin for these vesicles (Harischandra *et al*., 2018). A more focused investigation of the fluid found in these structures may prove more illuminating, as would demonstration that putative IVM targets are similarly expressed in reproductive tissues of adult female *B. pahangi* and *A. suum* to account for the potent IVM activity we noted in those species. Li *et al*, (2014) also observed tissue-specific *avr-14* expression in embryonic stages within gravid females. This corroborates the findings of Moreno *et al*. (2014) who noted strong localization of *avr-14* around the ES pore of *B. malayi* mfs, earmarking this structure as another, perhaps more predictable, site of EV secretion in this stage that lacks reproductive tissues or a through gut.

Whether these EVs have their biogenesis in reproductive tissues, the excretory system or some other secretory route, the pathways by which IVM inhibits their secretion is obscure and will require a more thorough description of the microscopic anatomy of key tissues and a better understanding of IVM targets expressed therein. For example, despite the recognition that parasitic nematode ES systems secrete a complex suite of molecules believed to be essential for successful parasitism (Allen & Maizels, 2011; Hewitson *et al*., 2009; Hoerauf *et al*., 2005; van Riet *et al*., 2007), the ultrastructure and transcriptional topography of the ES pore region has largely been uninvestigated. The intersection between EVs as an important mechanism for host manipulation during infection, the inhibition of their secretion by IVM at therapeutically relevant concentrations and time frames, and the localization of putative IVM targets in critical stage-specific tissues, provides strong rationale for addressing this knowledge gap.

A significant outcome from the work presented here is the demonstration that EV secretion from adult female and mf stage filarial nematodes (the stages that one could argue are most relevant to LF control programs) can be quantified and the effect of extraneously applied compounds on this secretion measured. Using this assay, we detected an IVM-sensitive EV secretion phenotype that perhaps correlates better with therapeutically relevant IVM concentrations than does assaying parasite motility, and in our experience is a more convenient and reproducible assay than that used to measure protein secretion from these worms. It may also be a better predictor of IVM mode of action. If the therapeutic mechanism of action of IVM is to inhibit immunomodulatory protein secretion from mf parasites then albendazole, which has been reported to inhibit protein secretion from *B. malayi* mf faster and more comprehensively than IVM (Moreno *et al*., 2010) (although we did not observe this), should also be an effective microfilaricide. Albendazole, however, is ineffective against mf stage filarial nematodes (Critchley *et al*., 2005). This suggests inhibition of EV secretion may be a preferred characteristic of anti-filarial drugs and therefore assaying this phenotype would be of significance to future drug discovery efforts aimed at developing new anti-filarial compounds, certainly those that function like IVM. In its favor, EV quantification would provide a consistent screening assay that would be comparable across different species of parasites, however, EV quantification is not high-throughput and does require additional EV isolation steps and specialized equipment for EV visualization. Recent technological advances may provide platforms that could be leveraged to streamline EV quantification assays and overcome these drawbacks. For example, we have contributed to an on-chip microfluidic device that utilizes a label-free photonic crystal biosensor to detect and discriminate host EVs from those secreted by parasitic nematodes based on differential expression of EV surface markers (Wang *et al*., 2018). This type of platform combines minimal sample processing with high throughput potential and does not require EV labelling, overcoming the disadvantages of traditional EV quantification and could be leveraged in drug discovery efforts centered on EV secretion as an assay endpoint. Another potential use for this type of platform are the early detection of parasite infection. The use of EVs as a nematode diagnostic has been seeded by the focus on EVs as diagnostic markers for cancer detection. Current advanced technologies involve using surface enhanced Raman scattering (SERS) or localized surface plasmon resonance to detect tumor-derived EVs in body fluids (Mehmet *et al*., 2017; Thakur *et al*., 2017; Zong *et al*., 2016). In addition, the miRNA cargo of EVs has been of interest as biomarkers for various cancers (Kosaka *et al*., 2019). Similarly, a microfluidic on-chip device has potential to identify parasite EVs from host biofluids. Current efforts towards this goal are aimed at identifying secreted parasite markers that that could be incorporated in such a design, or used in more simple assay formats such as PCR (Quintana *et al*., 2017; Tritten *et al*., 2014). Finally, the assay we describe here may be an example of a relatively simple *in vitro* assay to test or validate the emergence of anthelmintic resistance. The Fecal Egg Count Reduction Test is the gold standard for detecting resistance to anthelmintics like IVM. Alternative *in vitro* assays complement FECRT and include hatching and development assays, molecular tests and, of course, motility assays (Kotze & Prichard, 2016). The EV secretion assay could be added to this list if it could reliably, and with sensitivity, detect resistance to drugs such as IVM in a standardized fashion. There is some evidence for this potential; we previously detected differences in IVM susceptibility based on EV secretion for two strains of canine heartworm, *D. immitis* (Harischandra *et al*., 2018).

Collectively, our data show that the secretion of EVs from different parasitic nematode species can be assayed and the effects of anthelmintic drugs or lead compounds on this physiological process can be measured. IVM consistently inhibited EV secretion against all species and most life stages investigated, with the exception of male worms; other anti-filarial drugs did not. These findings provide new insight into the stage-, sex- and species-specific pathways and pharmacological regulation of EV secretion in parasitic nematodes. The data is significant because, given the emerging immunomodulatory role of EVs at the host-parasite interface, it provides new evidence that the therapeutic mechanism of IVM, in part, involves inhibition of parasite EV secretion.

## Disclosure of Interest

The authors report no conflict of interest.

## Notes

This work was supported by the National Institutes of Health under Grant AI117204.

### Competing Interest Statement

The authors have declared no competing interest.

