## Supplemental Materials for "Ivermectin inhibits extracellular vesicle secretion from parasitic nematodes"

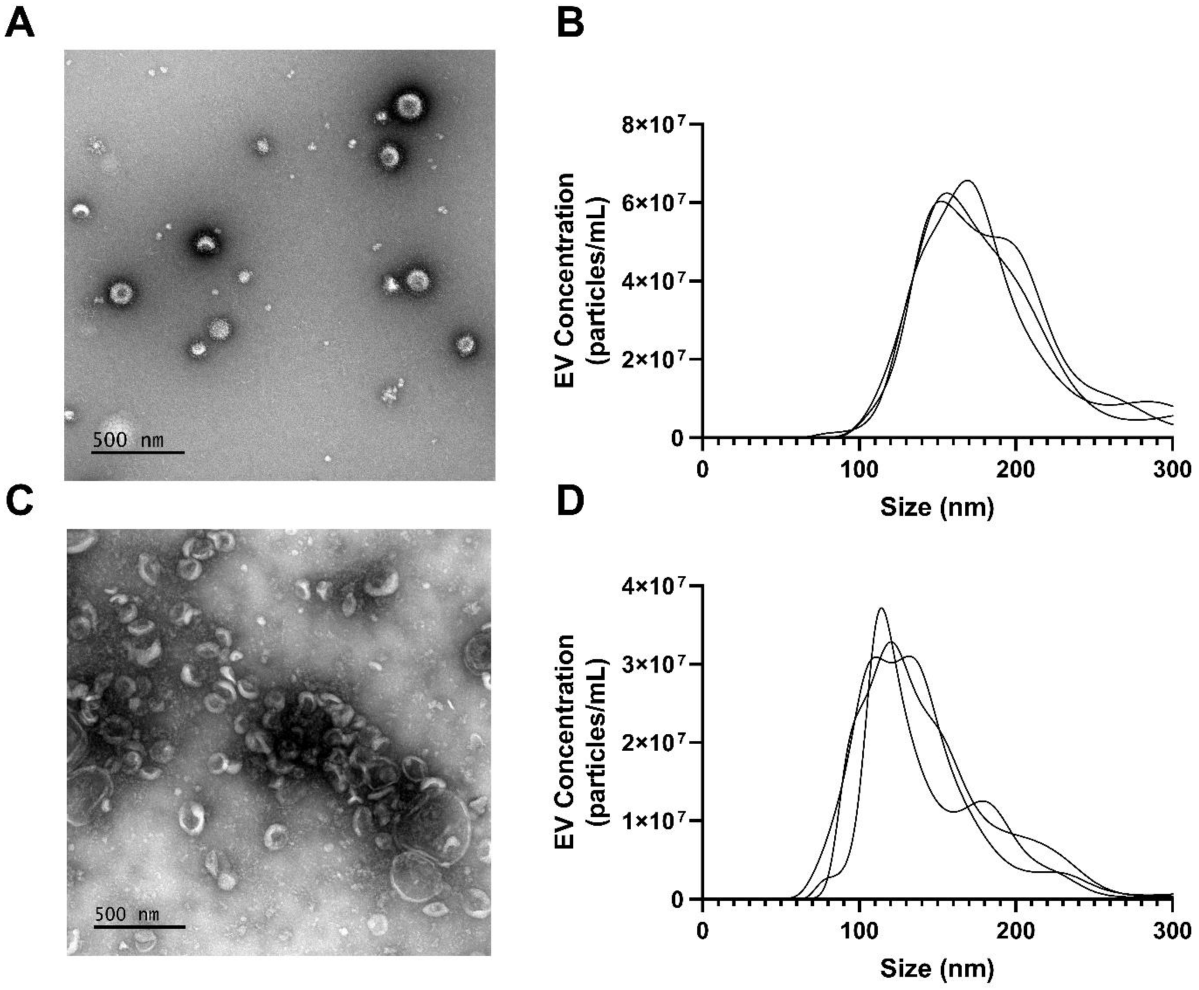


**Supplemental Figure 1. Validation of EVs via transmission electron microscopy (TEM) and NTA**

Adult Female *A. suum* or *B. malayi* mf were cultured and spent media was collected every 4 hrs for up to 72 hrs. EVs were isolated via differential ultracentrifugation as described. A two µl aliquot of purified EV preparation was placed onto a carbon film grid for 1 min. The drop was wicked to a thin film and two µl of uranyl acetate (2% w/v final concentration) was immediately applied for 30 sec., wicked, and allowed to dry. Images were taken using a 200kV JEOL 2100 scanning and transmission electron microscope (Japan Electron Optics Laboratories, LLC, Peabody, MA) with a Gatan OneView camera (Gatan, Inc. Pleasanton, CA). In addition, EVs were quantified using NTA. Representative electron micrographs of *A. suum* (A) and *B. malayi* mf (C) EVs. Representative NTA quantification trace of *A. suum* (B) and *B. malayi* mf (D) EVs.

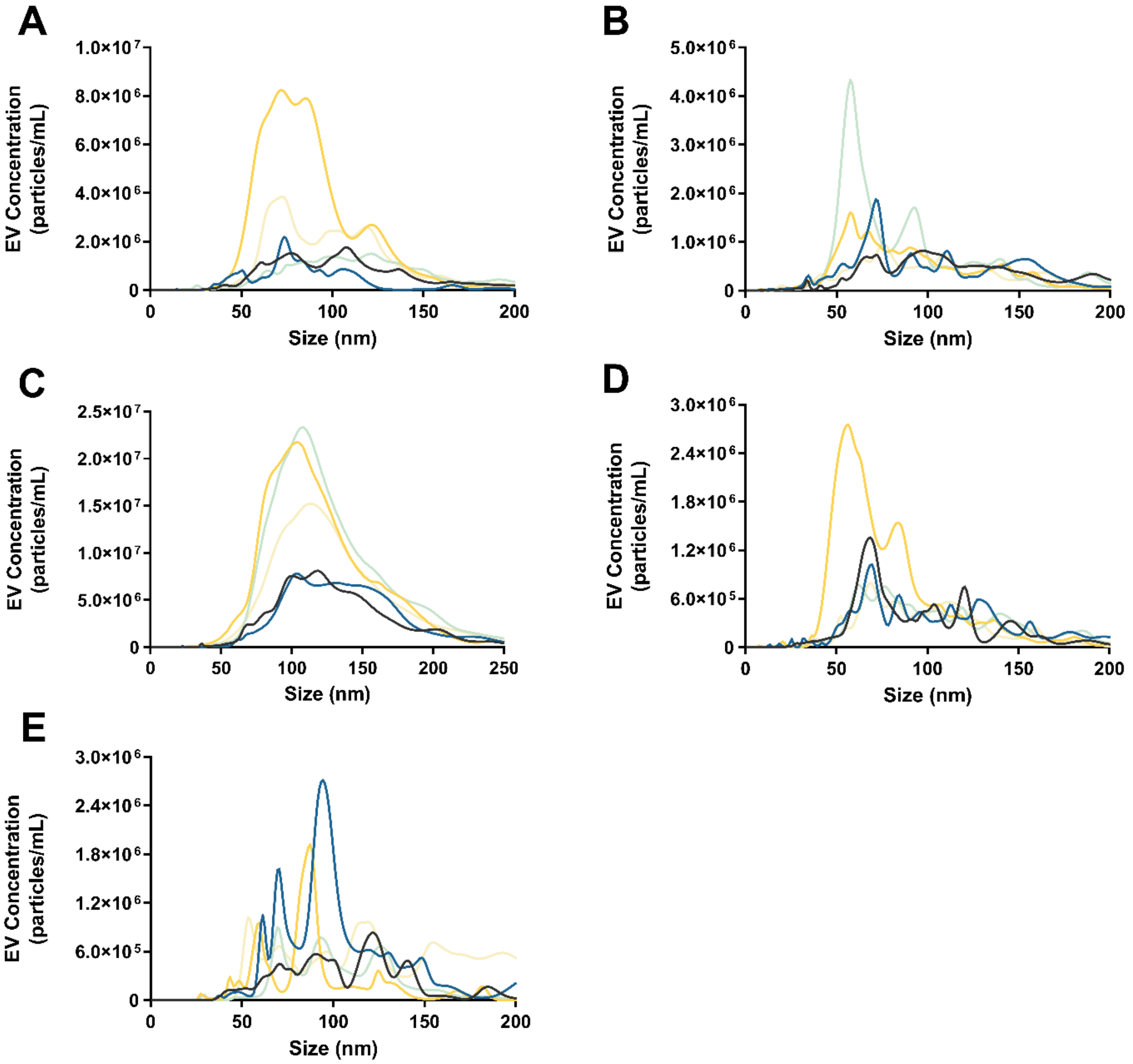


**Supplemental Figure 2. Anthelmintic treatment does not alter morphology of EVs secreted**

Control and drug treated parasites were cultured and spent media was collected after 24 hrs. EVs were quantified using NTA. Neither IVM, ABZ, DEC or LEV affected the physical characteristics of EVs secreted from *B. malayi* adult males (A), *B. malayi* L3 (B), *B. malayi* mf (C), *B. pahangi* adult male (D), or *B. pahangi* mf (E). N = 3 (minimum). ▬ = DMSO, ▬ = IVM, ▬ = ABZ, ▬ = DEC, ▬ = LEV.
